# Coat color allele and mtDNA haplotype distributions in Russian cat populations: a citizen-science research

**DOI:** 10.1101/165555

**Authors:** Olga O. Zaytseva, Yakov A. Tsepilov, Ekaterina S. Yudayeva, Artem Kasianov, Ivan V. Kulakovskiy, Maria Logacheva, Daria Belayeva, Anna A. Gracheva, Ksenia Homenko, Anastasia Knyazeva, Ekaterina Kolesnikova, Daria Nikolayeva, Mariana R. Bevova, Pavel M. Borodin, Yurii S. Aulchenko

**Affiliations:** Institute of Cytology and Genetics SD RAS, Novosibirsk, Russia, 630090; Novosibirsk State University, Novosibirsk, Russia, 630090; European Research Institute for the Biology of Ageing, University of Groningen, University Medical Center Groningen, 9713 AV Groningen, The Netherlands; Vavilov Institute of General Genetics, Russian Academy of Sciences, Moscow, Russia, 119991; Engelhardt Institute of Molecular Biology, Russian Academy of Sciences, Moscow, Russia, 119991; Moscow State University, Moscow, Russia, 119991; Summer School for Molecular and Theoretical Biology, Pushino, Russia, 142290

**Author notes:** These authors contributed to work equally (Olga O. Zaytseva, Yakov A. Tsepilov). Correspondence to: Yakov A. Tsepilov, Institute of Cytology and Genetics SB RAS, 630090 Novosibirsk, Russia.

**Keywords:** domestic cats, coat color, mitotypes, Russian populations

## Abstract

This work has started as a project for the summer school in molecular and theoretical biology. Efficient teaching and learning modern methods of molecular biology and population genetics in a two-week study course for high school students needs an attractive subject. We chose phylogeography of the domestic cat as the subject of our project because 1) everybody likes cats; 2) cats are polymorphic for several coat color mutations, which can be easily detected by street survey or in the photographs; 3) samples for DNA extraction are easy to collect without posing ethical and biosafety problems; 4) rich background information of geographical distribution of coat color alleles and variation in mtDNA is available; 5) phylogeography of cat population in Russia is poorly studied and therefore new data collected in this large territory may shed a light on the global cat distribution and on the origin of the fancy breeds.

During the project students studied coat color and mitotype distribution across Russian random bred cats. The basics of field (observations of natural cat populations, hair collection), formal (inheritance of coat colors), mathematical population (allele frequency distributions and their comparisons, building phylogenetic trees, multidimensional scaling), and molecular (DNA extraction and amplification, quality control of DNA sequencing data) genetics were covered.

We scored coat color phenotypes in 1182 cats and sequenced mtDNA control region from hair samples of 38 cats from 18 geographical sites. Analysis of coat color alleles frequencies and mitotype distribution confirmed relative homogeneity of gene pools of Russian cat populations, indicating their recent origin. We found several unique mitotypes and demonstrated that OL1 mitotype, previously found only in Siberian fancy breed, was present in random bred cats from several Russian cities. This contributes to the discussion on the origin of the Siberian breed of cats, and supports the view that Siberians is a recent breed created in the 1980s by breeding of selected representatives of the random bred population.

## Introduction

For some obscure reasons cats attract so much attention of humans that anything connected to cats has an extremely high memetic fitness [1]. Thus, all other factors being equal the audience would much better understand scientific facts and methodology illustrated by cats rather than Drosophila. Fortunately, the domestic cat as a species is very polymorphic for easily visible Mendelian traits such as coat color and texture [2]. They provide textbook examples of genetic phenomena such as complete and incomplete dominance (non-agouti and piebald spot correspondingly), epistasis and pleiotropy (dominant white), X-linked inheritance and X chromosome inactivation (Orange). This makes domestic cat an ideal subject for teaching genetics [2,3].

Because of a high level of Mendelian variation and mostly random breeding, cat is a good model species for the studies in population genetics and gene geography. J.B.S.Holdane was the first scientist to notice this [4]. In 1940s he put forward a very simple but powerful research program which later resulted in hundreds of surveys of coat color allele frequencies in cat populations all over the globe [4] and finally in detailed genogeographic map and phylogeny of local cat populations [5–8]. The results of a relatively recent microsatellite-based study [9] were in a good agreement with the results of coat color based studies. All these studies demonstrated a subdivision of the domestic cat into four distinct phylogeographic groups: Asian, Mediterranean, West European, and East African. North American cats consistently grouped with cats from Western Europe, suggesting European settlers probably brought cats to the New World and the cat’s time in America has been too brief for significant genetic differentiation.

As Darwin pointed out in the Origin of Species “cats, from their nocturnal rambling habits, cannot be matched, and, although so much valued by women and children, we hardly ever see a distinct breed kept up; such breeds as we do sometimes see are almost always imported from some other country” [10]. This remains mostly true even now. Eighty five percent of fancy cat breeds recognized by international associations were established in the past 75 years on basis of so-called natural breeds (local forms) and many of them are based on single-gene variants. Molecular genetic analysis indicates rather close resemblance between modern breeds derived from the local forms and random bred populations at the localities of their origin [11].

The molecular population genetics of fancy breeds and random bred populations was investigated with use of micro-satellite markers [9], and, more recently, sequencing of the mitochondrial DNA [12–14]. The studies of cat mtDNA [13,15,16] generated a comprehensive database of mtDNA control region (CR) mitotypes. The database included data on 1394 cats, of which 1027 were random bred cats from 25 distinct world-wide populations. Analysis of the database revealed 12 universal mitotypes with a frequency of >1% in the combined population set. However, most of the 12 mitotypes can be collapsed into two major mitotypes, A and D, as others are different from these by three mutations at most.

Russia is an interesting area in terms of cat phylogeography. It borders with three of four defined phylogeographic area and it is a likely source of at least two breeds: Russian blue and Siberian. There were several surveys of coat color gene frequencies in Russian populations. They indicated a relative homogeneity of genetic profiles of the cat populations across whole country and their resemblance to Mediterranean profile. A substantial contribution of West European cat populations was detected in sea port populations of St. Petersburg, Vladivostok and Petropavlovsk-Kamchatsky and their neighbors [17–20].

However, many Russian sites, which are interesting from the point of view of phylogeography and cat and human population dynamics have not been examined yet. A variability of mtDNA haplotype of the Russian cats remains totally unknown.

In our project we aimed to characterize coat color allele frequency and mitotypes of the control region of mtDNA in random bred cats of several Russian cities. This study started as a two-week project at Summer School in Molecular and Theoretical Biology (https://dynastybioschooleng.wordpress.com/) organized by NGO Dynasty Foundation and held in Academic city of Pushchino, Russia in 2013. The school program was oriented towards high school students. It was designed to integrate participants into research process in laboratories alongside with professional scientists with a possibility of obtaining novel results of scientific value. We chose gene geography of the domestic cat as the study subject because coat color genotypes can be easily detected by street survey or from photographs. Additionally, samples for mtDNA extraction can be easily collected without posing ethical and biosafety problems. Samples were collected by the students in their home cities. The time frame at the School was such that the students were to complete the project in approximately two weeks. During the first week students discussed with the staff members the general outline and workflow of the project, they were shown how to use laboratory equipment, and were introduced to molecular biology methods, such as DNA isolation, PCR reactions and sample preparation for sequencing. Second week was reserved for data analysis and discussion of the results.

Here we describe our experience in teaching and learning of modern population genetics using cats as a model and scientific results we obtained.

## Materials and Methods

### Coat color survey

High school students were selected to participate in the School based on their motivation for scientific research and knowledge in molecular biology. For this project we established a group for Cat geneogeography in the VK.com social network and invited the participants of the School to carry out coat color phenotype survey in their home cities. A letter containing sampling instructions were sent to the students and the staff of the school (See Supplementary Note I). We asked participants to collect information on the coat color of at least 100 randomly chosen cats, meaning that they were supposed to score any cats they meet, not only those remarkable and/or cute. We suggested to ignore obvious fancy cats score only one cat of a family.

The following coat variations were identified:

1. Orange or cream (*OO/OY*) versus black, blue or agouti spots (*oo/oY*), heterozygous females (*Oo*) are tortoiseshell;
2. agouti (*A_*) versus non-agouti or self-black (*aa*);
3. striped or mackerel tabby (*T_*) versus blotched or marble tabby (*t^b^ t^b^*);
4. intense (*D_*) versus blue dilution (*dd*);
5. dominant white (*W_*) versus pigmented (*ww*);
6. white spotting (*S_*) versus non-spotted (*ss*)
7. short hair (*ll*) versus long hair (*L_*)

The genetic basis for these phenotypes has been described by Robinson et al. [21]. Allele frequencies were calculated according to Borodin et al. [17]. To facilitate the phenotype scoring we suggested students to use our version of The Field Guide to Cat Phenotypes (Supplementary Note I). The Guide is based on the manuscript typewritten, illustrated by Kodak prints and privately distributed by Neil B. Todd among his colleagues. We also advised the participants to take pictures of at least some of the scored cats and upload them to our database together with phenotype descriptions (http://vk.com/albums-55078174). Before the project started, we received 561 photographs with 342 descriptions from 18 cities.

Analysis of these data revealed two weak points of our project.

Firstly, the geographical distribution of the sample sites was not optimal to study global and local distribution of allele frequencies. Majority of the samples originated from Central European part of Russia, samples from other regions were underrepresented. This problem could not be solved within the framework of the School, as students were selected irrespective of their geographical location. However the problem has partially been solved after the School when one of the students (E. Yudaeva) surveyed the cat populations along the Volga river (including cities of Kazan, Balakovo, Saratov, Volgograd and Astrakhan). She also established her own surveyors network covering additional cities of Rostov-on-Don, Krasnodar, Baku and Vladivistok. Her effort doubled the sample size and resulted in total of 1182 photos. Sample size and allele frequency for each location is shown in Table 1 and on the shared Google map (see Online materials).

**Table 1.** Coat color allele frequencies of cats collected in this study.

| Location | N | Allele frequency |  |  |  |  |  |  |
| --- | --- | --- | --- | --- | --- | --- | --- | --- |
|  |  | *O* | *a* | *d* | *t<sup>b</sup>* | *S* | *W* | *l* |
| Astrakhan | 58 | 0.360 | 0.657 | 0.350 | 0.000 | 0.312 | 0.009 | 0.394 |
| Baku | 50 | 0.167 | 0.663 | 0.204 | 0.186 | 0.308 | 0.020 | 0.245 |
| Balakovo | 97 | 0.135 | 0.674 | 0.323 | 0.000 | 0.257 | 0.005 | 0.321 |
| Barabinsk | 21 | 0.225 | 0.527 | 0.387 | 0.316 | 0.258 | 0.024 | 0.655 |
| Chelyabinsk region* | 47 | 0.163 | 0.788 | 0.374 | 0.236 | 0.301 | 0.043 | 0.484 |
| Kazan | 66 | 0.208 | 0.632 | 0.277 | 0.000 | 0.216 | 0.008 | 0.331 |
| Kostroma | 130 | 0.230 | 0.620 | 0.412 | 0.158 | 0.316 | 0.023 | 0.421 |
| Krasnodar | 98 | 0.412 | 0.749 | 0.363 | 0.261 | 0.435 | 0.036 | 0.465 |
| Novosibirsk region** | 86 | 0.146 | 0.702 | 0.366 | 0.000 | 0.366 | 0.024 | 0.638 |
| Pskov | 15 | 0.071 | 0.784 | 0.463 | 0.447 | 0.537 | 0.034 | 0.516 |
| Rostov-on-Don | 65 | 0.308 | 0.629 | 0.224 | 0.243 | 0.355 | 0.039 | 0.430 |
| Saint-Petersburg | 55 | 0.213 | 0.692 | 0.385 | 0.311 | 0.267 | 0.009 | 0.447 |
| Saratov | 106 | 0.167 | 0.679 | 0.313 | 0.000 | 0.366 | 0.019 | 0.521 |
| Tomsk | 42 | 0.325 | 0.655 | 0.418 | 0.000 | 0.293 | 0.024 | 0.408 |
| Vladivostok | 82 | 0.260 | 0.655 | 0.161 | 0.297 | 0.465 | 0.031 | 0.428 |
| Volgograd | 164 | 0.120 | 0.487 | 0.192 | 0.000 | 0.136 | 0.006 | 0.259 |

\* Ozersk, Metlino
\*\*Berdsk, Iskitim, Krasnoobsk

Secondly, many coat phenotypes were identified wrongly. While comparing descriptions provided by the students with the uploaded photos we found that of 1182 phenotypes 122 (10.3%) were incorrect. There were several common mistakes worth mentioning here. A yellowish variation of agouti was often confused with tortoiseshell (17.5 % of all phenotyping errors). Lighter lower lip was often mistaken for a white spot (3.9%). Non-agouti (*aa*), white spotting (*S_*), blue dilution (*dd*) and long hair (*ll*) were often overlooked (9.1%, 12.3%, 6.5%% and 9.1% respectively). Some surveyors failed to notice phenotypes tabby (1.3% of errors) and some mistook calico or wild type pattern for tabby (3.3%). Some of the Siamese were not recognized or mistaken for orange (9.1%).

The problem of misdiagnoses was solved by careful examination of each photograph by experts (PMB and OZ). All wrong diagnoses were commented online, so the students could learn from their mistakes. The final data contain verified diagnoses. However, some photographs did not allow us to properly determine tabby phenotype as cat’s backs were not visible. These cases were excluded from further calculation of *t^b^* frequencies.

This part of the work lead us to two conclusions:

All future projects on coat color population survey need to be documented by photos. It should be emphasized that the cat pictures should include pattern on the cat’s back in order to identify tabby phenotype.

Some results of past cat population genetics studies, even those published in scientific journals, should be taken with caution, especially when one or two localities shows exceptional frequencies of the alleles prone to misdiagnosis.

### Hair sampling

We asked the students to collect hair samples of random-bred cats in their neighborhoods. Hairs had to be sampled from the rear of the back of cats closer to the rump, aligned, fastened with a scotch tape (leaving the follicles end out) and placed in a labeled envelope. However, some of the hair samples lacked follicles, so that we had to discard them, as a reliable protocol for DNA isolation from hairs without follicles is time-consuming and requires some special equipment and reagents we lacked in the settings of the School.

A total 136 cat hair samples from 14 cities were collected by 15 students and 3 staff members of the school.: From those: 56 samples originated from European part of Russia and 80 from Asian part (Urals and Western Siberia) (see shared Google map in Online materials). This part of our project also suffered from uneven geographical distribution of the samples: more than 50% (76) of samples were provided by three students from Sankt-Petersburgh and two students from Ozersk.

### MtDNA typing

Root tags were trimmed from hair samples and incubated for 10 min at 100 C in 1,5 ml Eppendorf tubes with 150 ul of digestion buffer (provided by Dr. Maxim Filipenko, ICBFM SB RAS, Novosibirsk), then centrifuged at 14,000 g for 5 min. Supernatant was then transferred into clean tubes and used as a source of DNA matrix for PCR. Primers JHmtF3—gatagtgcttaatcgtgc and JHmtR3—gtcctgtggaacaatagg (Genbank IDs: U20753 and NC_001700) [14] were used for amplification of the domestic cat mtDNA control region. The size of amplified product is 491 bp including the primers.

PCR was performed in final reaction volume of 20 ul. Each PCR contained 2 ul sample DNA solution, 0.2 mM each dNTP, 1x Taq buffer (10 mM TrisHCl, pH 8.9, 55 mM KCl, 0.05% Tween-20, 2.5mM MgCl2), 0.3 pmol forward primer, 0.3 pmol reverse primer and 0.08 ul Taq polymerase (provided by Dr. Maxim Filipenko, ICBFM SB RAS, Novosibirsk). The amplification profile was following: 1) 94 ℃ for 2 min 30 sec of initial denaturation; 2) 5 cycles of 94C for 30 sec denaturation, 62 C for 30 sec annealing, 70 C for 1 min 30 sec extension; 3) 35 cycles of 94C for 30 sec denaturation, 60 C for 30 sec annealing, 72 C for 1 min extension; 4) 72 C for 3 min final extension. Samples were held at 10C until further processing. PCR product size was checked by electrophoretic separation in 1% agarose gel.

Two different methods of PCR product purification and sequencing were used to teach students a variety of molecular biology techniques.

ExoSAP clean-up was performed for 17 samples. Each 20-ul PCR sample was mixed with 2 U exonuclease I and 0.5 U thermostable alkaline phosphatase, followed by incubation at 37 C for 40 min, then at 85 C for 20 min. The rest of our samples were purified from PCRs with QIAquick Gel Extraction Kit (QIAGEN) from 1% agarose gel according to the protocol provided by the manufacturer.

Twenty five samples were sequenced using Sanger technique. In each Sanger reaction 2 ul of purified PCR product were used as a template for sequencing using the BigDye Terminator version 3.1 Cycle Sequencing Kit. Sequencing was performed on Applied Biosystems 3130xl by “Alfaferment”, Moscow, Russia. Next generation sequencing (NGS) technique was applied to 16 samples using Illumina MiSeq platform (Illumina, USA). Nextera XT DNA sample prep kit (Illumina) were used for library preparation from purified PCR products. After PCR and purification the libraries were pooled in equimolar amounts and sequenced using MiSeq Reagent Kit v2 (Illumina) with 250bp read paired-end protocol. Two samples were sequenced with both methods, giving 43 samples in total. Five samples (four by Sanger and one by NGS) produced sequences of poor quality and were excluded from further analysis resulting in 38 successfully sequenced samples. All successfully sequenced samples were used for alignment and mitotype elucidation.

For some samples sequenced with NGS technology several different nucleotides were observed at a single position of read pileups covering the studied regions (see Supplementary Table 5). In particular, there were several positions with sequence coverage ≥100x exhibiting different alleles in significant number of cases (with the major allele frequency less than 90%). While such positions may point to heteroplasmy, it does not provide complete evidence due to genomic numts repeats possibly affecting PCR results or read mapping. Possible occurrence of heteroplasmy was discussed with the students, thus they were introduced to the concept of heteroplasmy in general, but, unfortunately, in the settings of the summer school we were not able to conduct additional experiments to prove or exclude heteroplasmy in our samples.

### Analysis of sequence data

Sanger sequencing data were assembled with BioEdit 7.2.0 (http://www.mbio.ncsu.edu/BioEdit/) and Staden Package Software 1.6.0 (http://staden.sourceforge.net/). Forward and reverse sequences were assembled into contigs, and consensus sequences were generated. Manual editing was used to resolve any discrepancies between forward and reverse sequences. If manual editing could not resolve minor discrepancies, the sample was not used in further analysis. NGS reads were demultiplexed according to the barcodes. Raw reads were preprocessed by removing extremely short reads (< 10 bp) and polyN-masking low quality regions. Resulting reads were aligned to cat genome assembly (UCSC felCat5) using Bowtie2 (http://genomebiology.com/2009/10/3/R25), see Supplementary Table 4 for details. The consensus sequences were extracted from the reads pileups covering the studying mtDNA region selecting the most frequent nucleotides in each position. For each sample the resulting consensus sequences were trimmed and aligned to the 402 bp minimal variation consensus sequence and variable nucleotide positions were identified. All sequences were aligned with published reference sequences [13] using BioEdit 7.2.0 software (http://www.mbio.ncsu.edu/BioEdit/). Distance matrix was generated and all mitotypes were elucidated according to the previously published datasets [13].

### Statistical and phylogenetic analysis

The program NETWORK 4.6 (Fluxus Technology Ltd., Clare, Suffolk, UK) was used to create a phylogenetic network, which incorporated all possible shortest least-complex maximum-parsimony phylogenetic trees from the set of generated mitotypes.

The table of coat allele frequencies was processed in *ade4* package [22]. Nei’s distance was calculated for alleles at *O, a, t^b^, S* loci for each pair of cities. After that 2D multidimensional scaling were performed and visualized using R software.

Comparison of resampling data (repeated sampling of the same cities in different years) was performed using Fishers exact test for the same loci. If there was a significant difference for at least one locus, the resampling was scored as discordant. The threshold p-value< 0.0003 was chosen using Bonferoni correction: 0.01/(Number of populations*Number of loci).

Table of mitotype distribution was created and merged with the mtDNA haplotypes published earlier [13]. Euclidean inter-population distance matrix was calculated only for the populations with more than 20 observations. Two distance matrices were calculated: one using the information on unique mitotypes (which were merged into single ‘others’ mitotype) and another ignoring it.

Phylogenetic trees were generated using “phangorn” R package [23] with implemented methods of hierarchical cluster analysis. The “upgma” function with average linkage clustering was used. Package “PCPS” was used to define groups (clades) in a phylogenetic tree according to the selected threshold of 0.8. “Pvclast” ackage was used for hierarchical clustering analysis with bootstrapping and calculating of phylogenetic trees.

## Results and Discussion

### Coat color allele frequency

Estimates of frequencies of the seven alleles scored in 16 localities within the framework of this project are shown in Table 1 and on-line cluster map (see Online materials).

We added our data to the database of published data on the cat populations of former USSR countries [17,18,20,24–29]. This database has been collected, verified and kindly provided for this study by Dr. S.K. Kholin from the Institute of Marine Biology, Far Eastern Branch of Russian Academy of Sciences. The pooled database contains the data on 9832 cats from 56 cities collected in period from 1977 to 2015 yy (see Supplementary Table 2).

After filtering by sample size (no less than 30 cats per city), we found 11 cities with repeated sampling (see Supplementary Table 6). Eight of them showed significant resampling differences in the allele frequencies (Fisher exact test after Bonferroni correction p < 0.0003). We suppose that these differences may be determined by the microgeographic variation of the cat populations within the same cities or/and differences in scoring methods (street observation, shelter or veterinary clinic data), rather than historical changes in gene frequencies. For further analyses, we used data collected at the latest time point (54 cities, 6979 cats).

Almost all populations showed a high frequency of non-agouti (*a*) and Piebald spotting (*S*) alleles. Blotched Tabby (*t^b^*), which alter the patterns of coat coloration, are rare or missing in the populations of Baku and Russian cities on the Volga river. In Siberia marble cats are extremely rare too. We observed a moderate frequency of this allele in the populations of Vladivostok, St. Petersburg and Rostov-on-Don. This might be due to the fact that these cities are important trade ports. Cats carrying *t^b^* allele might have been brought to these cities by the maritime routes from Western Europe (mainly from Britain) where tabby cats are frequent [29]. The non-agouti (*a*) and other darkening alleles as *s*, *o* and *t^b^* are regarded as alleles of urbanization (Clark, 1975) [30]. We haven’t detected a significant correlation between the human population number and the pooled frequency of these alleles (R²~0.01, p>0.05).

Using most reliably diagnosed alleles *t^b^, O, a, S* we calculated genetics distances between the populations (see Materials and Methods) and performed multidimensional scaling (Supplementary Figure S1). There was no clear pattern except weak cluster that could correspond to Volga river cities from the south (Astrakhan) to the north (Kostroma). Volga has always been the most important trade route, which apparently facilitated intensive gene flow between the populations along the river and a moderate isolation of these populations from those located beyond this corridor.

For further analysis, we chose the representative cities from Russia (sample size more than 200) and all other cities from other countries (in total 19 cities, 3635 cats) and performed the same analysis as for Russian cities (Supplementary Figure S2). Again, no clear pattern was detected, except Gomel and Pokrov (former Ordzhonikidze) were outliers.

Thus, analysis of coat color alleles frequencies revealed relative homogeneity of gene pools of Russian cat populations, indicating their recent origin.

### Common and unique mitotypes

Table 3 shows the frequency of mitotypes found among Russian cats compared to worldwide data. All sequences are shown in Supplementary Note II. In total we found 14 mitotypes in our set of 38 samples (Supplementary Table 3). Four major mitotypes (A,B,C,D) account for >65% of mitotype diversity in worldwide random bred population [13]. Three of them were also present in our samples: A (16 cases), B (3 cases) and D (5 cases). Mitotype C which was previously found in 15 out of 19 populations was not observed in our sample set. This is consistent with the previous studies, in which at least three major mitotypes were observed in all studied populations [13]. However, mitotypes B and D were absent in Ural samples (Supplementary Table 3), probably, due to small sample size from this area (n=12).

**Table 3.**
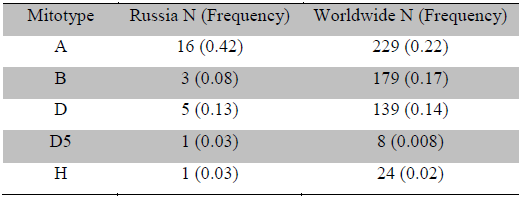
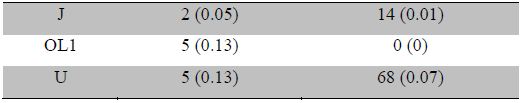
Mitotype occurrence in Russian cat populations compared with their occurrence in random bred worldwide cat populations ([13]).

| Mitotype | Russia N (Frequency) | Worldwide N (Frequency) |
| --- | --- | --- |
| A | 16 (0.42) | 229 (0.22) |
| B | 3 (0.08) | 179 (0.17) |
| D | 5 (0.13) | 139 (0.14) |
| D5 | 1 (0.03) | 8 (0.008) |
| H | 1 (0.03) | 24 (0.02) |
| J | 2 (0.05) | 14 (0.01) |
| OL1 | 5 (0.13) | 0 (0) |
| U | 5 (0.13) | 68 (0.07) |

In addition to three major mitotype we found three minor mitotypes: H (1 case), J (2 cases), U (5 cases), OL1 (5 cases) and D5, a derivative of a major mitotype D (1 case).

The already described mitotypes were observed in 33 of 38 cat examined. Each of five remaining cats had its unique mitotype, which have not been described before. Mitotype named *RUC_1* appears to be a derivative of mitotype A, *RUC_3* is a derivative of mitotype E, *RUC_4* - of mitotype I. *RUC_2* and *RUC_5* are outlier mitotypes, which are only distantly related to any major variants (see Figure 1).

**Figure 1.**
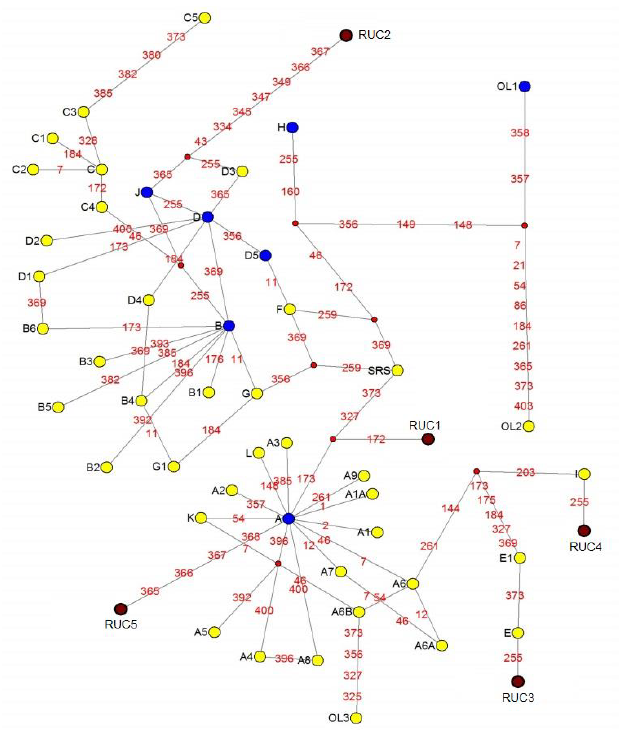
Network diagram of Russians cat mitotypes linked to the common mitotype network. Circles encompassing mitotypes are not to scale, although A is the most common. Subtypes are indicated at nodes. Small red dots indicate theoretical intermediary mitotypes predicted by network analysis. Blue nodes indicate mitotypes that were presented in our samples. Numbers on branches indicate nucleotide position changes in the sequence alignment with the Sylvester reference sequence (SRS). Three unattributed groups (OL1–3) and unique sequences (RUC1-5, dark red color) are indicated as well.

Interestingly, we found five cats with the mitotype OL1 (three from Urals and one from St. Petersburg and from Volgograd). This mitotype has not been previously detected in any random bred population [13], but has been found in 7 of 1394 fancy cats. Four carriers of this mitotype were found among 24 Siberian fancy cats examined. It was also found in one Birman cat, one Scottish Fold and one Turkish Van [13].

A relatively high frequency of OL1 among Russian random bred and Siberian fancy bred cats adds to the discussion on the origins of the Siberian Cat breed. One of us (PMB) witnessed an establishment of the Siberian breed in USSR. Two separate groups of cat breeders (one in St. Petersburg, another in Novosibirsk) decided to “resurrect” a legendary Siberian Cat, which in fact has never existed anywhere besides fairy tales and nursery rhymes. They established the standard of the breed based on the fairy tales: agouti long hair strong cat, with large, well rounded paws and large full tail, broad chest, broad foreheads. The founders of the breed were selected for a desired phenotype from random-bred and stray cats. The first standard of the breed was approved in 1990 by Soviet Felinological Federation. Within two years the breed standard was registered by World Cat Federation (WCF) and subsequently recognized by other international felinological organizations. Same story about the origin of Siberian cat is voiced by Mucha et al, 2011 [31].

The recent establishment of the Siberians from random bred Russian cats is reflected in higher levels of genetic variation detected in this breed. Lipinski et al. (2007) genotyped nineteen microsatellite markers in fancy and random bred cat populations from the USA and concluded that Siberian cats had variation comparable with a random-bred population [32]. High variation in Siberian cats was also noted by Kurushima et al. (2013) [11], who analyzed a panel of 38 short tandem repeats, 148 intergenic and five phenotypic single nucleotide polymorphisms in fancy cat breeds.

### Mitotype distribution worldwide and in Russian random bred population

Since our sample sizes from different locations were too small to detect population structure, we pooled them into two large groups: European and Asian (denoted as EuRUS and SURUS respectively). The Asian part included samples from Siberian and Ural towns: Ozersk, Ekaterinburg, Martianovo, Novosibirsk, Tomsk. The European part included samples from Mitotype distribution table and phylogenetic tree were created for these two populations (see Supplementary Figures S3-S4). Using euclidean distances between frequencies of common mitotypes by multidimensional scaling and hierarchical clustering method we did not detect a difference in mitotype distribution between these groups. Thus we observed little genetic diffreences between Russian European and Asian cat populations while comparing geographic distribution of mitotypes or coat colors. This homogeneity may reflect a very recent (100 - 200 years ago) establishment of the Asian populations and intensive gene flow from the European part of USSR during industrialization (1920-1930s) and massive evacuation during World War II (1940s).

We compared the mitotype distribution in pooled sample on the Russian cats with the published data on various populations of random bred cats (Figure 2 and Supplementary Figure S5). We detected a clustering of the populations divided into three phylogeographic groups: West Asia, Europe and former British colonies. Korean population was strong outlier due extremely high frequency (>60%) of E mitotype.

**Figure 2.**
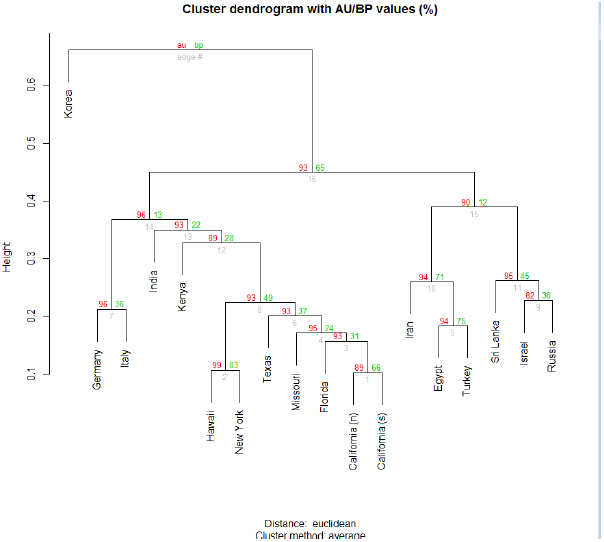
Hierarchical clustering phylogenetic tree of world random bred cat populations based on mitotype distribution. Phylogenetic trees were computed using hierarchical clustering method. The bootstrap method was used to calculate AU/BP values of the dendrogram (red and green numbers on graph respectively). AU (Approximately Unbiased) p-value computed by multiscale bootstrap resampling, which is more accurate than BP value as unbiased p-value. BP is a bootstrap probability. Gray number is a number of edge.

Russian cat population is close to those of Israel, Iran, Turkey, Egypt, Sri Lanka mostly due to relatively high frequency of mitotype D. Thus we can assume that Russian cat population belongs to West Asian group. Former British colonies characterized by relatively high frequencies of mitotypes B and C. Interestingly, all USA populations lay close to each other forming one big cluster (North America distinct group). Populations of Germany and Italy occupy an intermediate position between the populations of former British colonies American and West Asian populations.

## Conclusions

Our observation of high frequencies of OL1 mitotype in Siberia reinforces the hypothesis of recent (1980s) origin of the Siberian breed from Russian random bred and stray cats. Relative homogeneity of coat colors and mitotypes in the populations of cats from European and Asian parts of Russia is consistent with the history of recent large influx of humans from European to Asian part of Russia.

Population genetics of cats is a topic which allows teaching fundamentals of genetics to (high school) students in an entertaining and productive way. While we have asked students to do some background reading, most of the topics were covered on site within two-week summer school. These included elements of field (observational study of natural populations, hair sampling), formal (inheritance of coat colors), mathematical population (allele frequency distributions and their comparisons, building phylogenetic trees, multidimensional scaling), and molecular (DNA extraction and amplification, quality control of DNA sequencing data) genetics.

## Online materials

VK.com cats photo database together with their phenotype descriptions - http://vk.com/albums-55078174

Cats photo and hair samples distribution online map - https://www.google.com/maps/d/viewer?hl=en&authuser=0&mid=1I3zkhlq8hmiL28-42wIf4VTf7Zk

Coat color allele frequency online cluster map - https://www.google.com/maps/d/viewer?mid=1n7VUZ4dEQeqgTkJgLuOR8rt1VxQ&usp=sharing

## Acknowledgements

The authors thank the students, staff, and organizers of the School for Molecular and Theoretical Biology of the Dynasty Foundation (2013) for sample collection, fruitful environment and discussion. In particular, we thank Fyodor Kondrashev and Anya Piotrovskaya. Special thanks go to Ivan Bogachev, Regina Sharipova, and Alina Zakirova, who contributed more than 50% of the samples. We are also grateful to Maxim Filipenko for his advice on setting up the DNA isolation protocol and help with reagents.

## Authors contributions

YT, OZ, MB, YA, PB planned and supervised the study. EY, KA, GA, HK, KA, KE, ND, YT, OZ, YA collected data and contributed data for analysis. ML sequenced samples. EY, KA, GA, HK, KA, KE, ND, YT, KI, OZ, YA, MB performed experiments and data analysis and discussed and interpreted the results. YT, OZ, KI, YA, PM wrote the manuscript.

## Supplementary Information Legend

**Supplementary Note I –** instructions for sample collection

**Supplementary Note II –** FASTA sequences

**Supplementary Tables –** supplementary table 1-supplementary table 6

**Supplementary Figures –** supplementary figures S1-S6.

